# Refinement and reactivation of a taste-responsive hippocampal network

**DOI:** 10.1101/744714

**Authors:** Linnea E. Herzog, Donald B. Katz, Shantanu P. Jadhav

## Abstract

Animals need to remember the locations of nourishing and toxic food sources for survival, a fact that necessitates a mechanism for associating taste experiences with particular places. We have previously identified such responses within hippocampal place cells [1], the activity of which is thought to aid memory-guided behavior by forming a mental map of an animal’s environment that can be reshaped through experience [2–7]. It remains unknown, however, whether taste-responsiveness is intrinsic to a subset of place cells, or emerges as a result of experience that reorganizes spatial maps. Here, we recorded from neurons in the dorsal CA1 region of rats running for palatable tastes delivered *via* intra-oral cannulae at specific locations on a linear track. We identified a subset of taste-responsive cells that, even prior to taste exposure, had larger place fields than non-taste-responsive cells overlapping with stimulus delivery zones. Taste-responsive cells’ place fields then contracted, as a result of taste experience, leading to a stronger representation of stimulus delivery zones on the track. Taste-responsive units exhibited increased sharp-wave ripple co-activation during the taste delivery session and subsequent rest periods, which correlated with the degree of place field contraction. Our results reveal that novel taste experience evokes responses within a preconfigured network of taste-responsive hippocampal place cells with large fields, whose spatial representations are refined by sensory experience to signal areas of behavioral salience. This represents a possible mechanism by which animals identify and remember locations where ecologically relevant stimuli are found within their environment.

## Results and Discussion

To investigate how hippocampal place cells responded to tastes during running, we recorded the activity of CA1 neurons (*n* = 143, mean ± SEM: 28.6 ± 4.49 neurons/session; **Figure S1A**) as rats (*n* = 5) ran back and forth on a linear track during a novel taste experience (**Figure 1A**). Isolated units were classified as putative pyramidal cells (82.5%, 118/143) or interneurons (17.5%, 25/143; **Figure S1B**). Most pyramidal cells (99.2%, 117/118) could be categorized as place cells [1, 8] and exhibited spatially reliable firing over the course of a session (**Figure S1C**); a subset of these units (32.5%, *n* = 38/117) responded to tastes (**Figure 1B, Table S1**), consistent with previous findings [1]. Taste responses were observed only in cells whose place fields overlapped with delivery zones during the ‘Tastes’ session (**Figure 1C**), supporting previous work demonstrating that hippocampal sensory responses are gated by place cells’ spatial firing properties [1, 8, 9].

**Figure 1.**
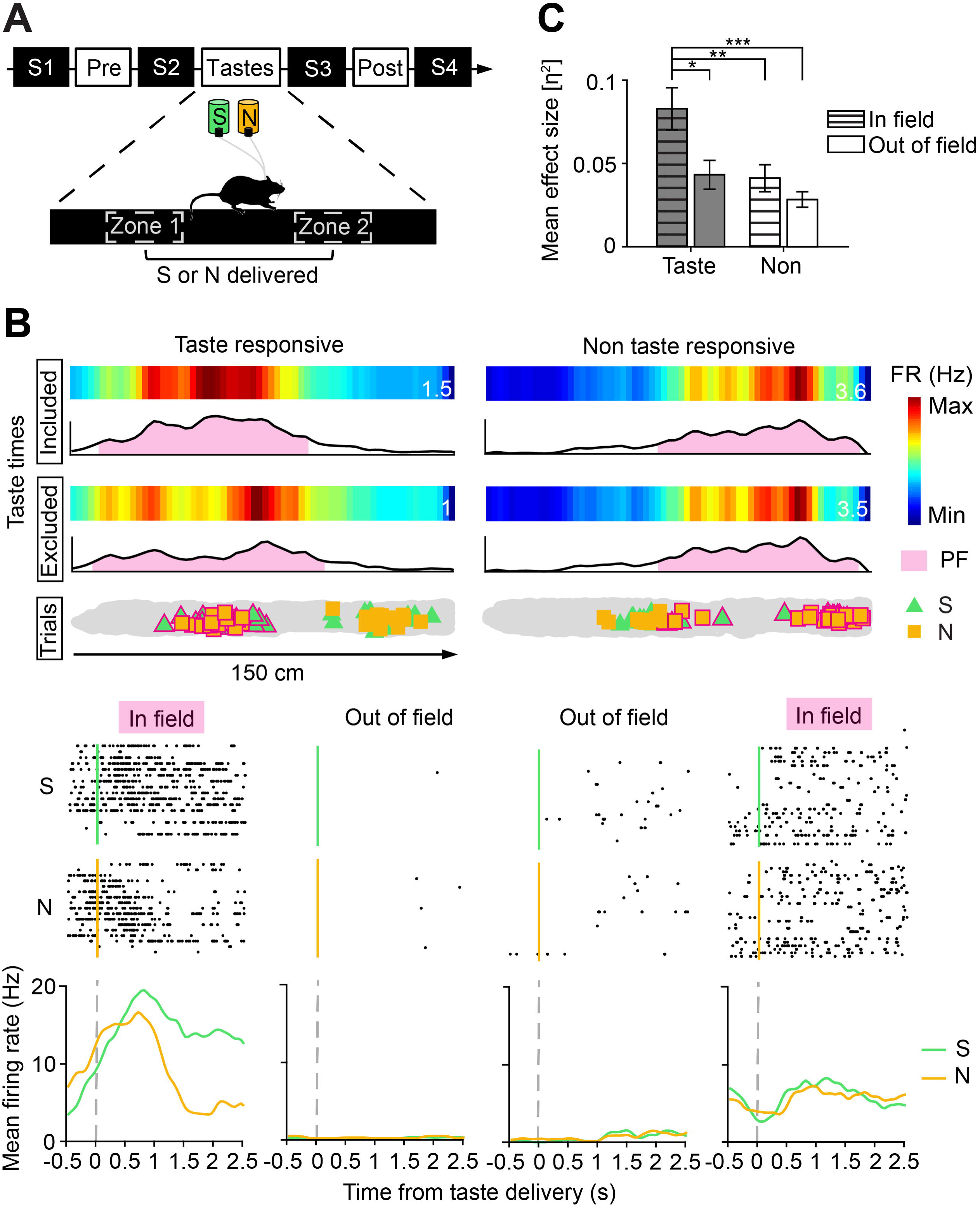
Experimental design and example taste responses. ***A***, Experimental schematic. Each experiment day consisted of seven sessions: three track-running sessions (Pre, Tastes and Post), interleaved with four sleep sessions (S1-4). For the Pre and Post probe sessions, rats ran on the track in the absence of tastes. During the Tastes session, rats received a randomized delivery of either sweet (green = S, 4 mM saccharin) or salty (yellow = N, 100 mM sodium chloride) taste solution upon entering either of the two designated regions on the track (Zone 1 and Zone 2). Neurons from the dorsal CA1 hippocampal region were recorded continuously throughout all sessions (see also **Figure S1** and **Table S1**). ***B***, Example in- and out-of-field taste responses. Top panels depict the firing rate (FR) maps (with peak FR denoted in Hz) and line plots for an example taste- and non-taste-responsive cell during the Tastes session, calculated with taste delivery periods either included or excluded from the analysis. Below, taste trial locations for the two example sessions are shown, with in-field trials (shown here using the taste-times-excluded place field) outlined in pink. Middle panels depict raster plots of each cell’s in- and out-of-field responses to each taste. Black dots indicate spike times during each individual trial aligned to the time of taste delivery. Bottom panels show the mean FR in response to saccharin or NaCl deliveries occurring in- and out-of-field for each cell. As previously reported [1], responses are only found within the taste-responsive cell’s place field. ***C***, Magnitude of taste responsiveness/specificity (ƞ^2^) for the in- and out-of-field regions of taste- (*n* = 20) and non-taste-responsive (*n* = 22) cells that met our criteria (≥ 10 trials in- and out-of-field). ƞ^2^ was highest (2-way ANOVA, cell type: \*\**p* = 0.0038, epoch: *\*\*p* = 0.0019, interaction: *p* = 0.13) for taste-responsive cells in-field (multiple comparisons tests, Taste/Out-of-field: \**p* = 0.012; Non/In-field: \*\**p* = 0.0094; Non/Out-of-field: \*\*\**p* = 2.1e-04). Similar results were obtained when taste delivery times were included in place map generation.

We previously established, using a passive taste delivery paradigm: 1) that a subset of hippocampal neurons encode taste palatability; 2) that these taste responses are gated by (i.e., only appear when the rat is in its) place fields; and 3) that taste-responsive neurons tend to have larger place fields than non-taste-responsive cells [1]. However, it remains unclear when these differences in spatial selectivity emerge, and how they evolve with taste experience. We therefore compared the place field sizes of taste- (*n* = 38) and non-taste-responsive (*n* = 79) place cells in the track-running session immediately preceding a novel taste experience (**Figure 2A**). Taste-responsive cells had larger place fields than non-taste-responsive cells before the rats’ first exposures to saccharin and NaCl solutions (**Figure 2B, S2A**), revealing a link between initial place field size and subsequent responsiveness to tastes (**Figure 2C, S2B**). However, we found no direct relationship between taste response magnitude and place field size (*n* = 42 cells, Pearson’s correlation: *r* = −0.23, permutation test, *p* = 0.14), arguing against the hypothesis that larger place fields simply connote higher sensitivity to responses. In addition, we have previously shown that hippocampal gustatory responses specifically code for taste palatability (rather than responding to any and all tastes) [1], rendering the possibility unlikely that spatially sparse neurons are more indiscriminately sensitive to sensory modulation, regardless of modality.

**Figure 2.**
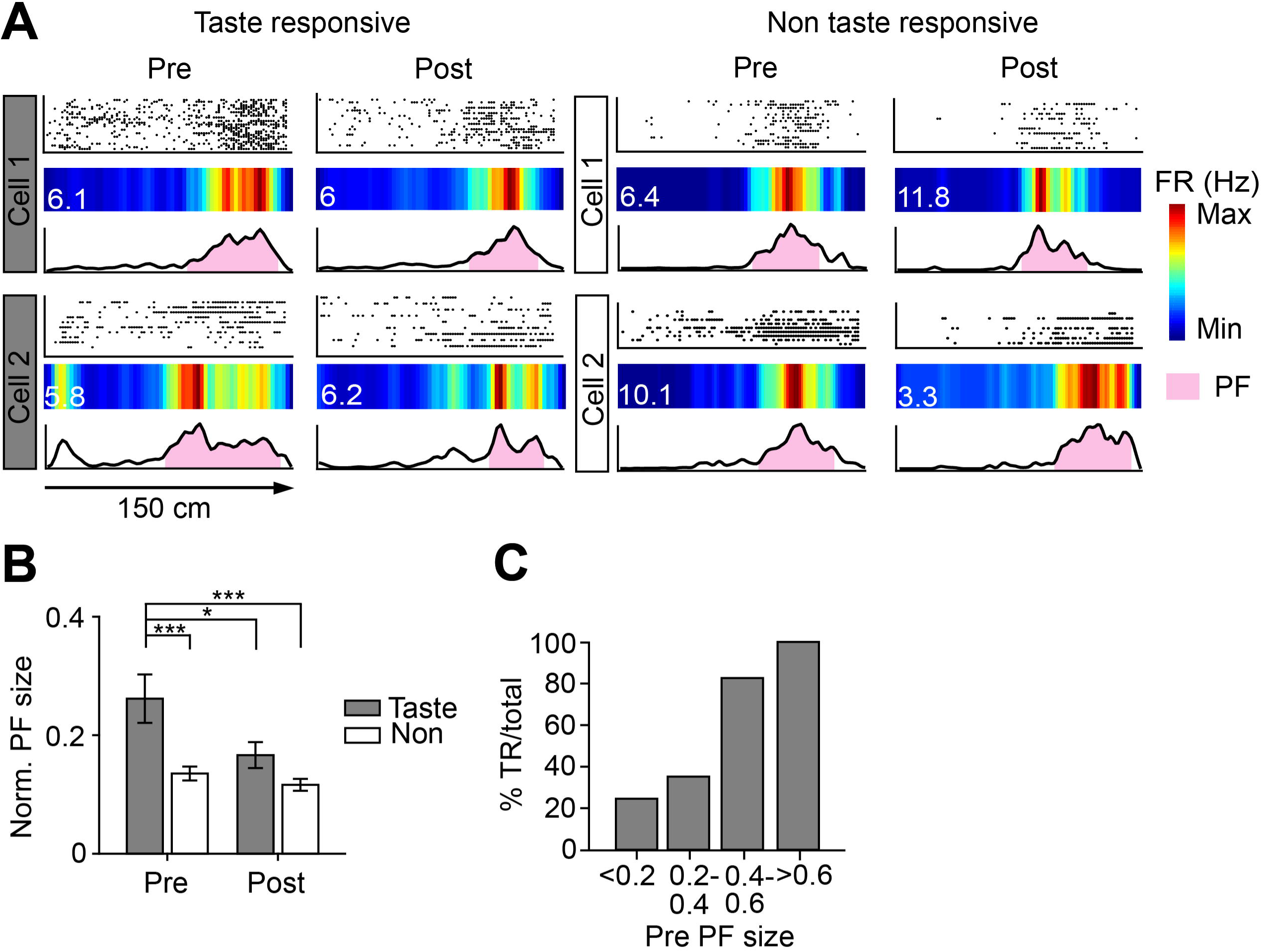
Taste-responsive place cells become spatially selective after taste experience. ***A***, Example taste- and non-taste-responsive place cell firing during the Pre and Post sessions. Top panels depict trial raster plots, with black dots indicating spike locations along each run of the track. Middle panels show the average linearized firing rate (FR) maps for each cell, with the peak FR denoted in Hz. Bottom panels depict FR maps as line plots, with place field (PF) boundaries indicated in pink. ***B***, Mean PF size for taste- (*n* = 38) and non-taste-responsive (*n* = 79) cells, normalized as a fraction of track coverage. Taste-responsive cells initially had larger PF (2-way ANOVA, cell type: \*\*\**p* = 9.8e-06, epoch: \*\**p =* 0.0037, interaction: *p* = 0.052), which contracted after taste experience (multiple comparisons tests, Pre/Non: \*\*\**p* = 4.6e-05; Post/Tastes: \**p* = 0.020; Post/Non: \*\*\**p* = 1.9e-06; see also **Figure S2**). ***C***, Likelihood of taste-responsiveness as a function of Pre PF size. Each bar represents the percentage of taste-responsive (TR) cells in each size bin, which increases with increasing PF size.

These results suggest that gustatory (or possibly reward) receptivity is to some degree a basic “feature” of the hippocampal system – perhaps reflecting differential input from the medial (MEC) and lateral (LEC) entorhinal cortices, which represent the two major cortical inputs to the hippocampus, encoding spatial and non-spatial information, respectively [10–13] — with initial differences in place field size potentially arising from preexisting synaptically connected ensembles [14–16].

Further, following taste exposure, taste-responsive cells’ place fields shrink, becoming comparable to those of non-taste-responsive place cells; non-taste-responsive cells’ place fields, meanwhile, did not change in size from the Pre to the Post session (**Figure 2B**). Analyses of local information content [17] yielded similar results. Since place fields are known to accumulate in rewarded locations [18–19], it is reasonable to hypothesize that this reorganization might function to localize place fields to taste delivery zones. However, we observed comparable degrees of location remapping in taste- and non-taste-responsive cells (**Figure 3A**) — no overall differences in Pre-Post place field location or place field-taste zone distance were found between groups (**Figure 3B**). Instead, taste-responsive cells exhibited high spatial overlap with taste delivery zones during the Pre session. Contraction of their place fields with taste experience decreased overall overlap (**Figure 3C**), but increased place field density in the delivery zones, such that the place field peaks of taste-responsive cells tended to more reliably overlap with taste zones in the Post session (**Figure 3D**). This refinement was observed in taste-responsive cells whose place fields initially overlapped with delivery zones (**Figure 3E**); lower proportions of non-taste-responsive cells exhibited overlap (29/38 vs. 44/79, X^2^ test, χ = 4.65, *p* = 0.031). These changes in spatial selectivity could not be attributed to interneuron firing, which tended to decrease during the Tastes session, then increase again during Post, becoming comparable to the Pre-session rate (1-way ANOVA, *p* = 3e-04, multiple comparisons tests, Pre vs. Taste: *p* = 2e-04, Tastes vs. Post: *p* = 0.032).

**Figure 3.**
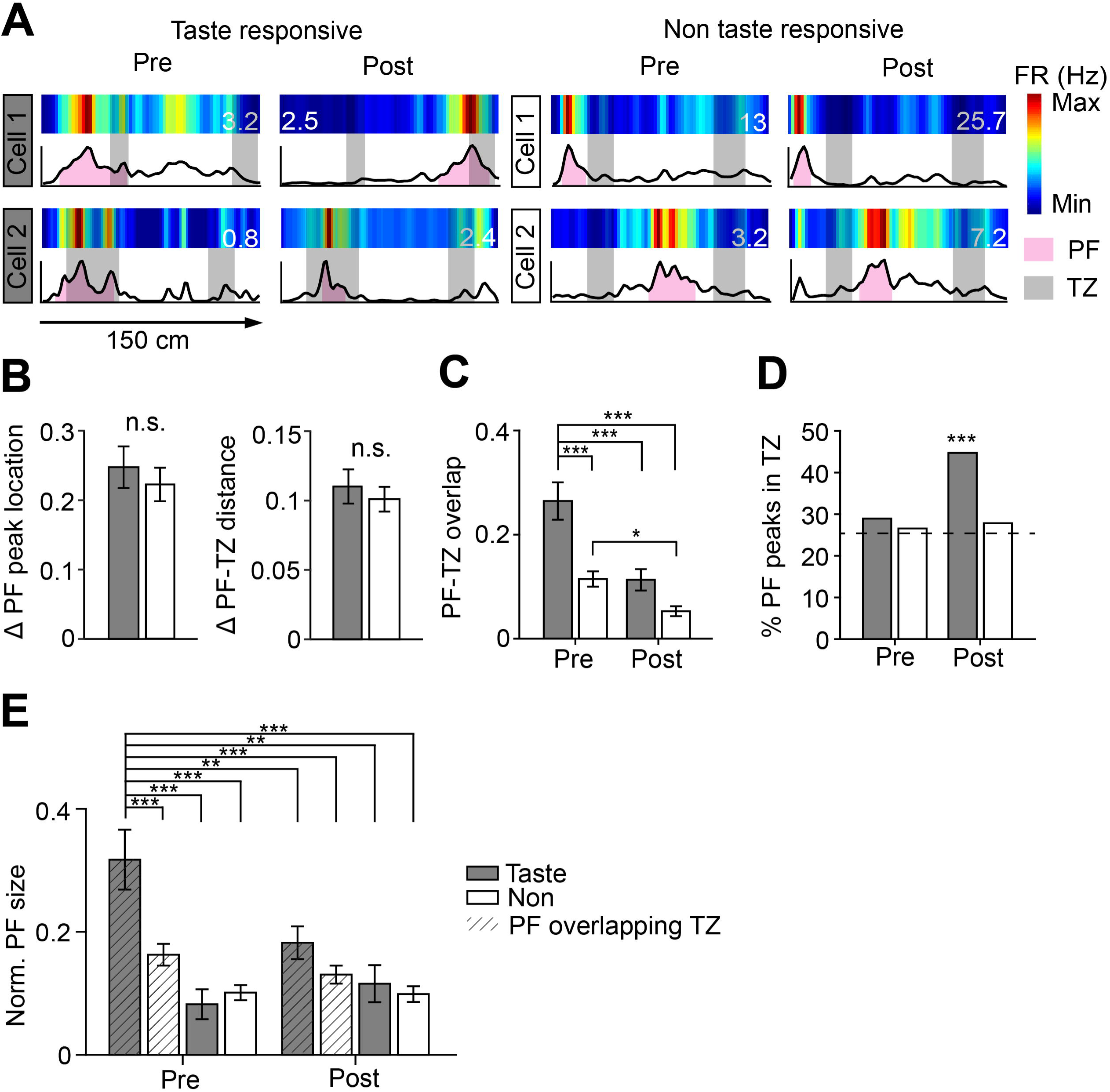
Taste-responsive cells’ place fields contract around areas of taste delivery. ***A***, Spatial firing maps of example taste- and non-taste-responsive cells during Pre and Post, with the peak firing rate (FR) denoted in Hz. Place field (PF) boundaries during each session are indicated by pink shading, while the peak distributions of trials (taste deliveries) during the Tastes session (taste zone, or TZ) are indicated by gray boxes. Note how taste-responsive cells’ PF overlap with TZ across taste experience, while non-taste-responsive cells’ do not. ***B***, Change in PF peak location and PF-TZ distance from Pre to Post. No differences in the mean Δ PF peak location (unpaired t-test, *p* = 0.54) or Δ PF-TZ distance (unpaired t-test, *p* = 0.55) were observed between groups or shuffled controls (paired t-tests, Δ PF peak location: Taste: *p* = 0.25, Non: *p* = 0.17; Δ PF-TZ distance, Taste: *p* = 0.89, Non: *p* = 0.31). ***C***, Mean PF-TZ overlap for taste- and non-taste-responsive cells. Taste-responsive cells’ PF exhibited high overlap with TZ during the Pre session (2-way ANOVA, cell type: \*\*\**p* = 9.6e-08, epoch: \*\*\**p* = 6.3e-08, interaction: \**p* = 0.020; multiple comparisons tests, Pre/Taste vs. Pre/Non: \*\*\**p* = 4.5e-07), which decreased with experience for both taste- (\*\*\**p* = 1.5e-05) and non-taste-responsive units (\**p* = 0.029). ***D***, Proportion of taste- and non-taste-responsive cells with PF peaks in TZ. Bars show the percentage of cells with PF peaks located within the TZ, compared to chance level (calculated based on average TZ coverage of the track). A greater-than-chance proportion of taste-responsive cells had PF peaks in the TZ during Post (binomial test, *\*\*\*p* = 3.9e-05), underscoring increased density of taste-responsive cell field peaks in TZ after experience. ***E***, Mean PF size as a function of PF-TZ overlap during Pre. TZ-overlapping taste-responsive cells had larger PF during Pre (3-way ANOVA, cell type: \**p* = 0.015, epoch: *p* = 0.10, overlap: \*\*\**p* = 3.7e-06, cell type x epoch: *p* = 0.42, cell type x overlap: \**p* = 0.013, epoch x overlap: \**p* = 0.018, cell type x epoch x overlap: *p* = 0.097) than any other group (multiple comparisons tests, vs. Pre/Non/Overlap: \*\*\**p* = 5.8e-05; Pre/Taste/No overlap: \*\*\**p* = 1.6e-04; Pre/Non/No overlap: \*\*\**p* = 1.5e-08; Post/Taste/Overlap: \*\**p* = 0.0039; Post/Non/Overlap: \*\*\**p* = 3.9e-07; Post/Taste/No overlap: \*\**p* = 0.0026; Post/Non/No overlap: \*\*\**p* = 1.0e-08), revealing that the contraction effect shown in Figure 2B is confined to taste-responsive cells with PF initially overlapping the TZ.

In summary: taste-responsive cells initially have large place fields that overlap with future locations of taste delivery as taste responses are gated by place fields; these place fields then contract following a novel taste experience, leading to a more precise representation of taste delivery locations in animals’ mental maps of the track. Non-taste-responsive cells do not exhibit this effect, instead representing the track continuously throughout taste experience.

One possible mechanism underlying this representational reorganization is the co-firing of goal-related assembly patterns during sharp-wave ripples (SWRs; [20–21]), a high-frequency hippocampal oscillation that is necessary for memory consolidation during wake [22] and sleep [23]. During ripples, the activity of sequences of place cells is typically “replayed” in a time-compressed manner [24–25]; sensory cues are known to influence this SWR reactivation [26] and replay content [27]. We therefore hypothesized that taste-responsive place cells would exhibit particularly strong SWR co-activation during waking behavior and sleep, as reorganization occurred.

To test this hypothesis, we identified SWR co-activation events (**Figure S3A**) in taste- and non-taste-responsive cell pairs (**Figure S3B**) whose place fields exhibited high and low spatial overlap. As expected [24, 28–30], highly overlapping cell pairs co-fired more during wake and sleep SWRs than pairs with low overlap (**Figure 4A**). This effect persisted when we calculated the mean co-activation differences for all possible combinations of cell pairs (Taste/Taste or Non/Non – Taste/Non) across sleep and wake (**Figure 4B**). Highly overlapping taste-responsive cell pairs exhibited higher co-activation differences than similarly overlapping non-taste-responsive cell pairs during taste experience and post-experience sleep (**Figure 4B, S4C**). This effect parallels the changes observed in place field size and spatial coverage on the track—we saw that taste-responsive cells’ place fields contract over areas of taste delivery following the Tastes session, while non-taste-responsive cells remain stable across taste experience. Furthermore, we observed a significant correlation between place field contraction (from Pre to Post) and SWR reactivation probability during the Tastes session (**Figure 4C, S4D**). These data support the hypothesis that reactivation during SWRs can help stabilize animals’ mental maps as they encounter tastes within their environment, with taste-responsive units coming to more precisely signify areas where food can be found.

**Figure 4.**
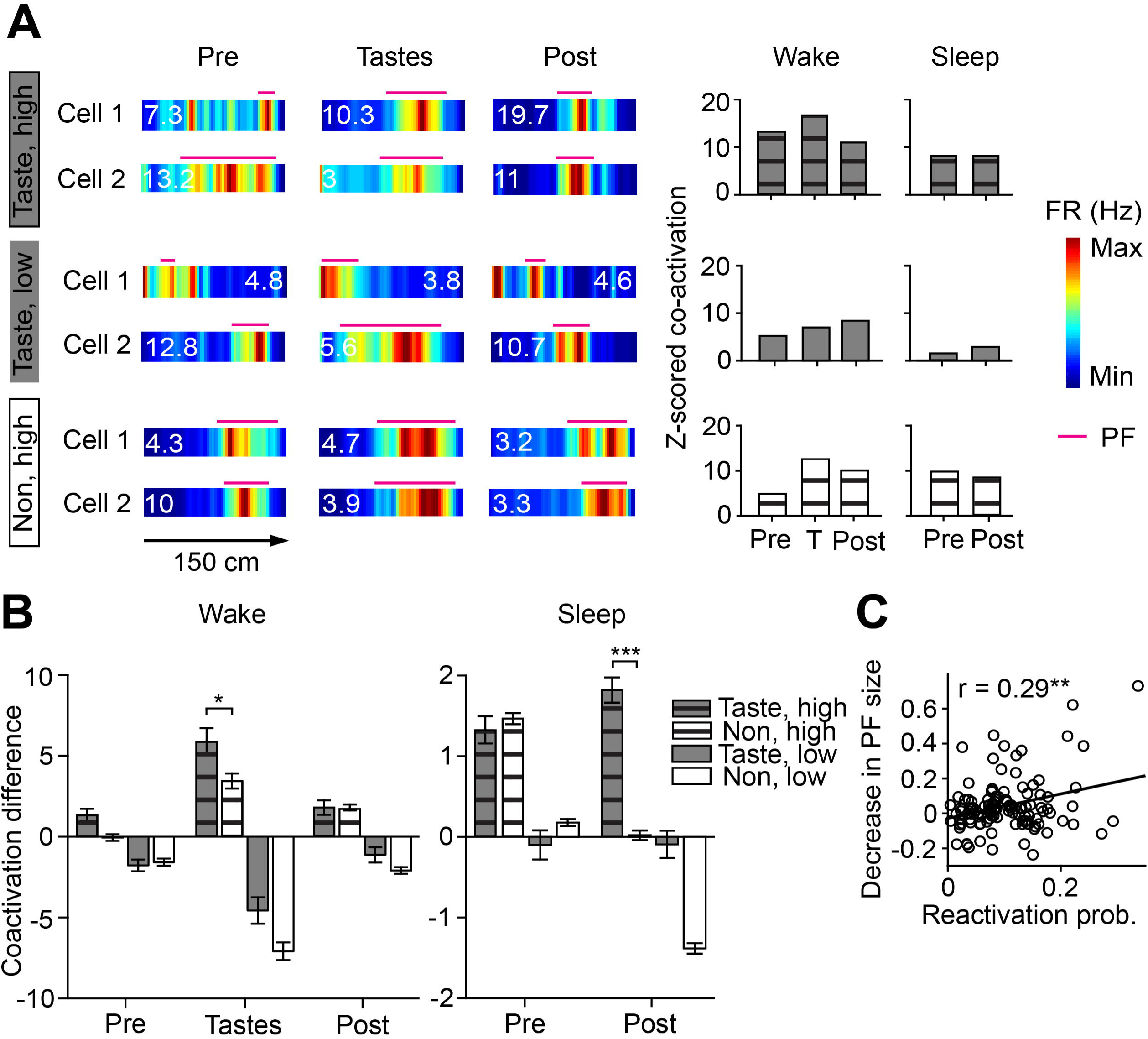
Taste-responsive cells exhibit increased SWR co-activation during taste experience. ***A***, Spatial firing and SWR co-activation for three example cell pairs (taste-responsive pairs with high and low overlap, non-taste-responsive pair with high overlap; see also **Figure S3**). Numbers on each place map plot denote peak spatial firing rate (FR) in Hz. Place field (PF) boundaries during each session are indicated by pink lines. The bar plots on the right show the Z-scored co-activation for each cell pair during wake and sleep SWRs. ***B***, co-activation differences between taste- and non-taste-responsive cell pairs across taste experience. Bars display the mean difference in SWR co-activation between same (Taste/Taste or Non/Non) and different (Taste/Non) cell pairs for all significant co-activation events. Significant differences were found for both wake (3-way ANOVA, cell type: \*\*\**p* = 3.7e-04, overlap: \*\*\**p* = 9.1e-74, cell type x epoch: \*\**p* = 0.0067, epoch x overlap: \*\*\**p* = 1.1e-33) and sleep (cell type: \*\*\**p* = 5.6e-09, epoch: \*\*\**p* = 4.6e-08, overlap: \*\*\**p* = 2.9e-39, cell type x epoch: \*\*\**p* = 2.3e-14). Post hoc comparisons revealed that taste-responsive cell pairs with high overlap had higher co-activation differences than non-taste-responsive cell pairs during taste experience (\**p* = 0.046) and post-experience sleep (\*\*\**p* = 1.1e-15). ***C***, Reactivation probability during the Tastes session is correlated with the decrease in PF size from the Pre to Post session, (*n =* 117 cells, Pearson’s correlation: *r* = 0.29; permutation test: \*\**p* = 0.002)

Overall, our data show that taste responsivity is intrinsic to a subset of hippocampal place cells, which initially possess larger place fields than non-taste-responsive cells prior to taste exposure. The introduction of novel tastes in a familiar environment led to the contraction of taste-responsive cells’ place fields over delivery zones—a refinement of animals’ internal representations of the track to emphasize locations of taste consumption. This sharpening of place fields only took place during animals’ first exposure to sweet and salty tastes (place field size, second experimental day: 2-way ANOVA, cell type: *p* = 1, epoch: *p* = 0.57, interaction: *p* = 0.96), revealing that the introduction of novel tastes in distinct spatial locations is what drives response plasticity, not unlike how place fields themselves classically emerge and stabilize over animals’ first exposure to a new environment [31].

The idea that rewarding or aversive experiences can reorganize hippocampal representations of an environment to encode points of behavioral salience is not new [18-19, 32-33], but we have shown that these shifts arise from changes in the spatial properties of a preexisting network of sensory-responding cells, which refine their spatial representations. One possible mechanism underlying this shift is the increased SWR co-activation of taste-responsive units with high spatial overlap, which correlated with the degree of place field contraction that we observed over taste experience. Our findings are consistent with previous observations that introducing novel stimuli, whether rewarding [18–19] or aversive [32–33], leads to the increased reactivation of behaviorally relevant trajectories during SWRs and reorganizes animals’ hippocampal representations to map out areas of high salience within an environment. These results provide further evidence for a flexible memory framework where hippocampal neurons can multiplex contextual information and sensory cues to update animals’ mental maps of an environment [4, 34–37].

## Acknowledgments

This work was supported by the Alfred P. Sloan Foundation (Sloan Research Fellowship in Neuroscience to S.P.J.), the Whitehall Foundation (S.P.J.), and the National Institutes in Health (NIH Grants R01 DC006666 and R01 DC007703 to D.B.K. and Training Grant T90 DA032435 to L.E.H.). We would like to thank Wenbo Tang for providing the rat illustration in the Graphical Abstract (available on Scidraw.io).

## Author Contributions

L.E.H. performed all experiments and analyzed the data. D.B.K. and S.P.J. supervised all aspects of the work. All authors conceived and designed the study and wrote the paper.

## STAR Methods

### Lead Contact and Materials Availability

Further information and requests for resources and reagents should be directed to the Lead Contact, Dr. Shantanu P. Jadhav (shantanu{at}brandeis.edu). This study did not generate any new reagents.

### Experimental Model and Subject Details

Five adult male Long-Evans rats (450-550 g, 4-6 months, Charles River Laboratories: RRID: RGD_2308852) served as subjects in this study. Rats were individually housed and kept on a 12 h light/dark cycle, with all sessions taking place around the same time during the light period. All surgical and experimental procedures were conducted in accordance with the National Institutes of Health guidelines and approved by the Brandeis University Institutional Animal Care and Use Committee.

### Method Details

#### Experimental design

After several weeks of habituation to daily handling, animals were pre-trained to seek water rewards delivered via reward well at either end of an elevated linear track, and habituated to an elevated, opaque sleep box [22]. After animals learned to run back and forth on the track, they were chronically implanted with a multi-tetrode drive and an intra-oral cannula (see *Surgical implantation and electrophysiology*).

Following recovery from the implantation surgery (∼7-8 days), rats were water-deprived to 85-90% of their *ad libitum* weight and re-trained using the same paradigm as described above, but with one change – water rewards delivered directly via intra-oral cannula instead of at reward wells. At ∼14 d after implantation, we performed daily recording sessions in which rats ran back and forth on the linear track during a novel taste experience (see **Figure 1A** and *Zoned taste paradigm*), where animals received sweet and salty tastes at two locations, located ∼1/3 and 2/3 of the way down the track. Palatable taste solutions, which naïve rats prefer over water [38–39], were used in order to motivate the animals’ behavior. Animals were naïve to these tastes prior to experiment day, as well as taste exposure in these locations, which were distinct from those experienced during pre-training. Only data from animals’ first day of exposure to this paradigm were used for the analyses described in the paper, unless otherwise specified. Following the conclusion of these zoned taste experiments, additional recordings were conducted while rats received a battery of basic tastes; data from these experiments has been presented previously to demonstrate that hippocampal neurons respond to tastes [1].

Following the conclusion of all experiments, we made electrolytic lesions through each electrode tip to mark recording locations. After 12-24 hours, animals were euthanized and intracardially perfused with 4% formaldehyde using approved procedures. Brains were fixed for 24-48 hours, cryoprotected (30% sucrose in 4% formaldehyde), and stored at 4°C. Brains were sectioned into 50 µm slices and stained with cresyl violet to confirm electrode placement in the hippocampal cell layer (**Figure S1A**).

#### Zoned taste paradigm

Each recording session typically lasted between 2 and 3 hours, and consisted of three running sessions on a 150 cm elevated linear track interleaved with four 15-20 minute sleep sessions in a ∼30 x 30 x 40 cm black box (rest box). The first (Pre) and last (Post) sessions on the linear track consisted of 15-20 minute periods in which animals ran back and forth on the track in the absence of tastes (mean number of track runs per session: 17.2 ± 3.7 during Pre, 14.0 ± 3.2 during Post). During the 30-90 minute Tastes session, rats received a pseudo-randomized delivery of either a sweet [4 mM saccharin (S)] or salty [100 mM sodium chloride (N)] taste solution at designated areas (“taste zones,” or TZ) located approximately 1/3 and 2/3 of the way down the track (mean number of taste trials per session: 91.2 ± 17.7). Rats consistently slowed down after taste delivery (average speed in the 2.5 s before taste delivery: 11.8 +/− 0.32 cm/s, average speed in the 2.5 s after taste delivery: 3.11 +/− 0.10 cm/s; unpaired t-test, *p* = 1.1e-122). The rat’s position was detected in real-time with an overhead monochrome CCD camera (30 fps) and LEDs affixed to the recording headstage, and used to trigger taste deliveries when animals entered into one of the two taste delivery zones. Taste solutions were delivered directly onto the tongue in ∼40 µL aliquots via two polyamide tubes inserted into the intra-oral cannula, with a separate tube for each solution to prevent the mixing of tastes. Animals had access to an additional 15-20 mL of water in their home cage after the end of each experiment day.

#### Surgical implantation and electrophysiology

Following pre-training on the linear track and habituation to the sleep box, animals were implanted with a microdrive array consisting of 25-30 independently moveable tetrodes in the right dorsal hippocampal region CA1 (−3.6 mm AP, 2.2 mm ML), and an intra-oral cannula. Each intra-oral cannula consisted of a polyethylene tube inserted beneath the left temporalis muscle and terminating anterolateral to the first maxillary molar, allowing for the precise delivery of taste solutions onto the rat’s tongue [38–40].

Electrophysiological recordings were conducted using a SpikeGadgets system [41]. Spikes were sampled at 30 kHz and bandpass filtered between 600 Hz and 6 kHz. Local field potentials (LFPs) were sampled at 1.5 kHz and bandpass filtered between 0.5 and 400 Hz. Over ∼14 d following surgery, tetrodes were gradually advanced to the CA1 hippocampal cell layer, as identified by characteristic EEG patterns (SWRs; theta rhythm) as previously described [22, 41–42]. Tetrodes were readjusted after each day’s recordings. Each animal had one hippocampal reference tetrode in corpus callosum, which was also referenced to a ground screw installed in the skull overlying cerebellum.

#### Unit inclusion

Spikes were sorted as described previously [1, 41–42]. In brief, single units were isolated offline based on peak amplitude and principal components (Matclust, M.P. Karlsson). Only well-isolated units with stable waveforms that fired at least 100 spikes per session were included in our analysis. Putative interneurons (*Int*) were identified on the basis of firing rate (> 8.5 Hz) and spike width (< 0.35 ms) parameters (**Figure S1B**). All other isolated units were classified as putative pyramidal cells (*Pyr*). We isolated a total of 143 neurons from five rats, conducted across five experiments, and recorded throughout the experiments. **Table S1** shows the distribution of cells across all five animals.

### Quantification and Statistical Analysis

#### Sharp-wave ripple detection

SWRs were detected as previously described [41–42] using the ripple-band (150-250 Hz) filtering of LFPs from multiple tetrodes. A Hilbert transform was used to determine the envelope of band-passed LFPs, and events that exceeded a threshold (mean + 3 SD) were detected. SWR events were defined as the times around initially detected events when the envelope exceeded the mean. SWR periods were excluded from place field analysis, similar to previous studies [1, 41–42].

#### Taste response properties

Hippocampal responses to tastes were characterized as described previously ([1]; for characterization of response properties in other parts of the taste system, see [40, 43–46]. Since we have already extensively described the coding content of taste responsiveness in these neurons [1], and given our use of only two tastes in the current context, here we perform (except when noted, see *Taste selectivity* below) a simpler test of taste responsiveness. In brief, neurons were classified as taste-responsive if they exhibited responses to taste presence (divergence from baseline) and/or identity (taste-specific differences in evoked firing) in the 2500 ms of spiking activity following each taste delivery; all other units were classified as non-taste-responsive (see **Figure 1B** for examples).

#### Taste selectivity

The magnitude of taste responsiveness and specificity for each cell was quantified using eta-squared (η^2^), a standard measure of ANOVA effect sizes that describes the proportion of variance in a dependent variable explained by each factor:

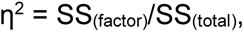

where SS is the sum of squares [1, 47]. In our analysis, we used the summed SS of the two main factors (time + taste) to calculate η^2^. As described below (see *In-field vs. out-of-field analysis*), a 2-way ANOVA was used to assess differences in η^2^ for the in-field and out-of-field regions of taste-responsive and non-taste-responsive place cells that fit our analysis criteria (> 1 Hz place field peak, ≥ 10 in- and out-of-field trials; **Figure 1C**). To determine if there was a general relationship between taste response magnitude and place field size, we used permutation tests to compare the Pearson’s correlation between in-field η^2^ and PF size during the Tastes session (*n* = 42 cells that fit our analysis criteria with taste delivery times excluded in place map generation, see *In-field vs. out-of-field analysis*) with 1000 cell ID-shuffled controls.

#### Spatial maps

To visualize the spatial firing properties of neurons, 2D position data were converted to linear positions, and one-dimensional occupancy-normalized firing rate maps (**Figure 1B, 2A, 3A, 4A**) were constructed using 2 cm bins and smoothed with a 1D Gaussian (σ = 5 cm) as previously described [22, 41]. Place fields from each session (‘Pre,’ ‘Tastes,’ and ‘Post’) were detected individually. Data from SWR periods (see *Sharp-wave ripple detection*) and periods of immobility (< 3 cm/s) were excluded from spatial map analysis.

Peak rates for each cell were defined as the maximum firing rate across all spatial bins in the spatial map. Each cell’s place field was defined as the largest group of neighboring spatial bins which had firing rates ≥ 20% of the peak rate (in some cases, multiple regions on the track met this criteria, in which case only the largest is depicted as the place field); sizes were then calculated by multiplying the number of bins by the bin size [48]. Only place cells (*n* = 117, defined as pyramidal cells whose unsmoothed peak firing rate exceeded 1 Hz [1, 8]) were included in analyses of spatial firing.

Within-session spatial reliability (**Figure S1C**) was measured by dividing each of the ‘Pre’, ‘Tastes’ and ‘Post’ sessions into three segments. The correlation coefficients of firing maps generated in each segment (first vs. middle, middle vs. last, first vs. last) were compared separately for taste-responsive (*n* = 38) and non-taste-responsive (*n* = 79) cells using 1-way ANOVAs [49]. Permutation tests using 1000 firing-rate-shuffled controls were used to assess the spatial reliability of individual neurons during the Pre session, and the proportions of spatially reliable taste- and non-taste-responsive cells were compared using a X^2^ test.

To compare place field sizes across sessions, place field size was normalized as a fraction of the track. A 2-way ANOVA was used to determine whether the mean place field size differed significantly between taste-responsive and non-taste-responsive place cells before and after animals’ first exposure to the zoned tastes paradigm (**Figure 2B**; this analysis was also conducted separately for repeated exposures; *n* = 6 taste-responsive and *n* = 49 non-taste-responsive cells across 4 sessions). Positional information at peak firing locations [17] was similarly analyzed across sessions and cell types. Importantly, these comparisons of place field size and local information content were between the Pre and Post sessions, which were identical in nature (neither involves taste delivery) and speeds maintained by rats traversing the track. The mean PF size for individual animals is depicted in **Figure S2A**. The percentages of taste- and non-taste-responsive cells in each spatial bin (0.2*n as a fraction of track coverage) were compared using X^2^ tests (**Figure S2B**). A plot of the percentage of taste-responsive cells in each spatial bin was also constructed (**Figure 2C**).

To determine how taste experience affects place field position (**Figure 3B**), the mean change in place field peak location (Δ PF peak location) and place field-taste zone distance (Δ PF-TZ distance) from Pre to Post were compared for taste- and non-taste-responsive cells using unpaired t-tests. The taste zones were defined separately for each experiment day using a Gaussian fit to determine the peak distribution of trial locations during the Tastes session (defined as μ ± σ for the normal distribution of all trial locations, calculated separately for each delivery zone).The closest taste zone to a place field’s peak location was used to calculate PF-TZ distance. Paired t-tests were used to compare both the mean Δ PF peak location and Δ PF-TZ distance for each group with controls that were calculated by randomly shuffling values from the Post session.

To compare the PF-TZ overlap across sessions (**Figure 3C**), the track was divided into 100 spatial bins, and we calculated which bins each PF and TZ spanned along the normalized track. PF-TZ overlap was defined as the number of taste zone locations that overlapped with a cell’s place field, divided by the total number of taste zone locations. 2-way ANOVAs were used to compare the PF-TZ overlap between taste- and non-taste-responsive cells before and after taste experience.

To determine if greater-than-chance numbers of cells in any group had place field peaks inside the taste delivery zone (**Figure 3D**), binomial tests were used to compare the observed vs. expected proportions of taste-responsive and non-taste-responsive cells with PF peaks in the TZ during Pre and Post. The expected proportion (25.4%) was calculated based on the average TZ coverage of the track, operating under the assumption that place fields exhibit equal coverage of the track.

To determine how initial overlap with taste delivery zones affects place fields (**Figure 3E**), taste- and non-taste responsive cells were grouped according to their TZ overlap during the Pre session (*n* = 29 overlapping taste-responsive cells, *n* = 9 non-overlapping taste-responsive cells, *n* = 44 overlapping non-taste-responsive cells, *n* = 35 non-overlapping non-taste-responsive cells); place field sizes were then compared between all groups using a 3-way ANOVA. The proportions of TZ-overlapping units were also compared using a X^2^ test.

#### In-field vs. out-of-field analysis

To analyze how cells responded to tastes delivered inside or outside of their place fields (**Figure 1B**), only place cells that contained at least ten in- and out-of-field deliveries of each taste were considered. This analysis was conducted separately using spatial maps calculated with taste delivery times (from 500 ms before to 2500 ms after taste delivery) included (*n* = 22 taste-responsive and *n* = 21 non-taste-responsive cells) or excluded (*n* = 20 taste-responsive and *n* = 22 non-taste-responsive cells). A 2-way ANOVA was used to assess differences between the mean in-field and out-of-field taste response magnitude (calculated using η^2^, see *Taste selectivity*) of taste-responsive and non-taste-responsive cells (**Figure 1C**).

#### Analysis of interneuron firing rates

To determine whether changes in place cell firing were influenced by interneuron activity in the taste zone, a 1-way ANOVA was used to compare the average normalized firing rates of interneurons in the TZ across sessions [50].

#### SWR co-firing

SWRs co-activation events were defined as ripples during which both cells in a given cell pair fired spikes (example in **Figure S3A**; locations of all awake ripples and co-firing events are plotted in **Figure S3B**). Co-activation of cell pairs during awake and sleep SWRs was calculated as previously described [30]:

If *n_A_* and *n_B_* represent instances when neurons A and B were independently active during N events, then the expected number of events during which both neurons were active (*n_AB_*) follows a hypergeometric distribution [51] with mean

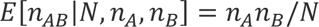

and variance

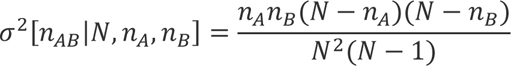

We use a Z-score to normalize across cell pairs with different activity levels (see right panels of Figure 4A):

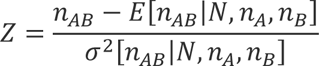

Only cell pairs that fired at least 100 spikes per session, had a peak firing rate exceeding 3 Hz during run sessions, and co-fired at least 5 times during SWRs were considered for analysis. Place field overlap for pairs of cells was defined as twice the sum of the overlapping areas of the linear rate curves divided by the sum of the areas of each curve [22]; all cell pairs with overlap ≥ 0.5 were classified as high overlap (see examples in top and bottom panels of **Figure 4A**), while all cell pairs with overlap < 0.5 were classified as low overlap (see example in middle panel of **Figure 4A**). For sleep SWRs, the closest running session was used to calculate spatial overlap.

The mean co-activation difference was defined as the difference in Z-scored co-activation between all combinations of same (Taste/Taste or Non/Non) – different (Taste/Non) cell pairs during awake running sessions (Pre, Tastes and Post) as well as the Pre rest session before the run sessions and the Post rest sessions after taste experience (averaged across the two rest sessions, S3 and S4). For these comparisons, same cell pairs (Taste or Non) with a greater spatial overlap than their corresponding different cell pair were classified as high overlap (High), while same cell pairs with a smaller spatial overlap than their corresponding different cell pair were classified as low overlap (Low). A 3-way ANOVA was used to compare the cell type, overlap and epoch effects on the mean co-activation difference during awake and sleep SWRs (**Figure 4B**; number of same-different cell pair combinations; wake: *n =* 96 Pre/Taste/High, *n* = 332 Pre/Non/High, *n* = 96 Pre/Taste/Low, *n* = 310 Pre/Non/Low, *n* = 74 Tastes/Taste/High, *n* = 320 Tastes/Non/High, *n* = 118 Tastes/Taste/Low, *n =* 322 Tastes/Non/Low, *n =* 107 Post/Taste/High, *n* = 305 Post/Non/High, *n* = 85 Post/Taste/Low, *n* = 337 Post/Non/Low; sleep: *n* = 261 Pre/Taste/High, *n =* 2221 Pre/Non/High, *n* = 182 Pre/Taste/Low, *n* = 4475 Pre/Non/Low, *n* = 248 Post/Taste/High, *n* = 3990 Post/Non/High, *n* = 195 Post/Taste/Low, *n* = 2706 Post/Non/Low). Means for individual animals are shown in **Figure S3C**.

In order to assess the relationship between SWR activation and place field refinement (**Figure 4C**), the reactivation probability was computed for all units as previously described [19]; a Pearson’s correlation between reactivation probability and the difference in PF size from the Pre to Post sessions was then calculated. Permutation tests using 1000 cell ID-shuffled controls were used to assess the significance of this correlation. Means for individual animals are depicted in **Figure S3D**.

#### Statistical analysis

All statistical tests were performed using custom routines in MATLAB (MathWorks, Natick, MA, RRID: SCR_001622) and evaluated at a level of α = 0.05, with a Bonferroni correction applied for multiple comparisons. Unless otherwise noted, values and error bars in the text denote means ± SEM.

### Data and Code Availability

The data and code used in this study are available upon reasonable request by contacting the Lead Contact, Dr. Shantanu P. Jadhav (shantanu{at}brandeis.edu).

## Supplemental Information titles and legends

**Figure S1. Histology, unit classification and spatial reliability, related to Figure 1.**

***A***, Histological verification of tetrode locations in intermediate dorsal CA1. ***B***, Classification of putative interneurons (Int, gray crosses) from pyramidal cells (Pyr, black circles) based on spike width (> 8.5 Hz) and firing rate (< 0.35 ms) parameters. ***C***, Spatial reliability of place cells across sessions. There were no significant changes in the mean correlation coefficient of taste-responsive (*n* = 38) and non-taste-responsive (*n* = 79) cells’ place field maps across each third of the session, indicating that spatial firing was reliable within each session (1-way ANOVA, Taste responsive, Pre: *p* = 0.99, Tastes: *p* = 0.21, Post: *p* = 0.55; Non-taste-responsive, Pre: *p* = 0.27, Tastes: *p* = 0.72, Post: *p* = 0.98). Single-cell analyses confirmed that similar proportions of taste- and non-taste-responsive cells exhibited spatially reliable firing during the Pre session (X^2^ test, X = 1.26, p = 0.26).

**Figure S2. Taste responsiveness is related to place field size during the Pre session, related to Figure 2.**

***A***, Place field size in Figure 2B shown for individual animals. ***B***, Histogram of taste-responsive and non-taste-responsive place field sizes. Stacked bars represent the percentage of cells within each cell type that fell within a given size bin (n*0.2 as a fraction of the track). Cells with smaller place fields were more likely to be non-taste-responsive (PF size < 0.2, X^2^ test, χ = 6.5, \**p* = 0.011), while cells with larger place fields were more likely to be taste-responsive (PF size = 0.4-0.6, χ = 7.5, \*\**p* = 0.0063; PF size > 0.6, χ = 6.4, \**p* = 0.011).

**Figure S3. SWR co-activation and distribution of significant events along the track, related to Figure 4.**

***A***, Example of one co-activation event following a taste trial, with saccharin delivery denoted by the vertical green line. Spikes from two taste-responsive cells are shown in black, with spikes that occurred during the same SWR inside the purple bar. SWRs were detected using the simultaneously recorded EEG filtered at 150-250 Hz (black traces above raster plot). The top trace shows the SWR in higher magnification. ***B***, Probability of SWRs and significant co-activation events for taste- and non-taste-responsive cell pairs at each location on the track during awake running sessions. For visualization, taste zones (TZ, gray shaded areas) are depicted as an average across sessions. ***C***, Mean co-activation differences from Figure 4B shown for individual animals (each column corresponds to a bar on the main plot). For each epoch, only animals with sufficient numbers of cell pairs are shown. ***D***, Mean reactivation probability vs. decrease in place field (PF) size from Figure 4C shown for individual animals.

**Table S1. Cell distribution across animals, related to Figure 1.**

Summary of the number of taste-responsive and total CA1 cells recorded from each animal. Only cells meeting the inclusion criteria (see STAR Methods) are reported. Putative pyramidal cells (Pyr) and interneurons (Int) were identified on the basis of firing rate and spike width parameters. Neurons were classified as “taste-responsive” if they exhibited responses to taste presence or identity.

