## Supplemental Information for "Refinement and reactivation of a taste-responsive hippocampal network"

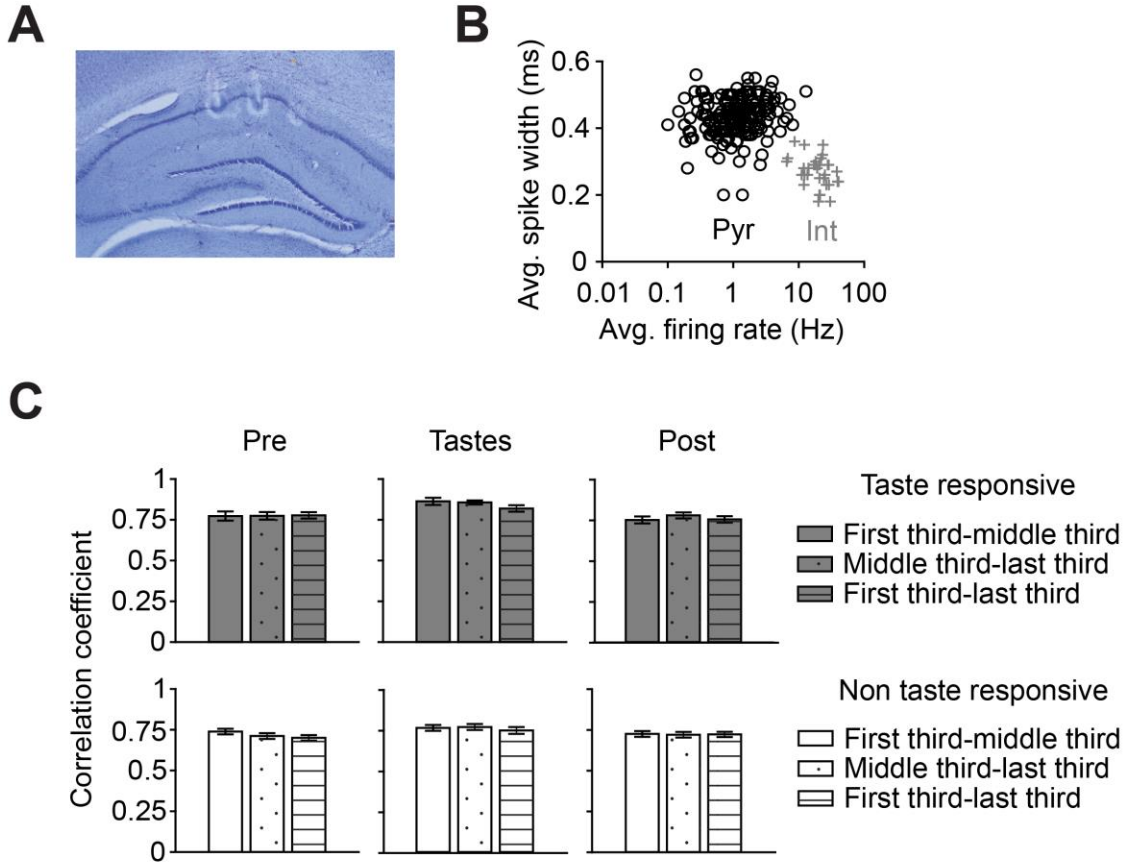


**Figure S1. Histology, unit classification, and spatial reliability, related to Figure 1.**

***A***, Histological verification of tetrode locations in intermediate dorsal CA1. ***B***, Classification of putative interneurons (Int, gray crosses) from pyramidal cells (Pyr, black circles) based on spike width (> 8.5 Hz) and firing rate (< 0.35 ms) parameters. ***C***, Spatial reliability of place cells across sessions. There were no significant changes in the mean correlation coefficient of taste-responsive ($n$ = 38) and non-taste-responsive ($n$ = 79) cells' place field maps across each third of the session, indicating that spatial firing was reliable within each session (1-way ANOVA, Taste responsive, Pre: $p = 0.99$, Tastes: $p = 0.21$, Post: $p = 0.55$; Non-taste-responsive, Pre: $p = 0.27$, Tastes: $p = 0.72$, Post: $p = 0.98$). Single-cell analyses confirmed that similar proportions of taste- and non-taste-responsive cells exhibited spatially reliable firing during the Pre session ($\chi^2$ test, $\chi = 1.26$, $p = 0.26$).

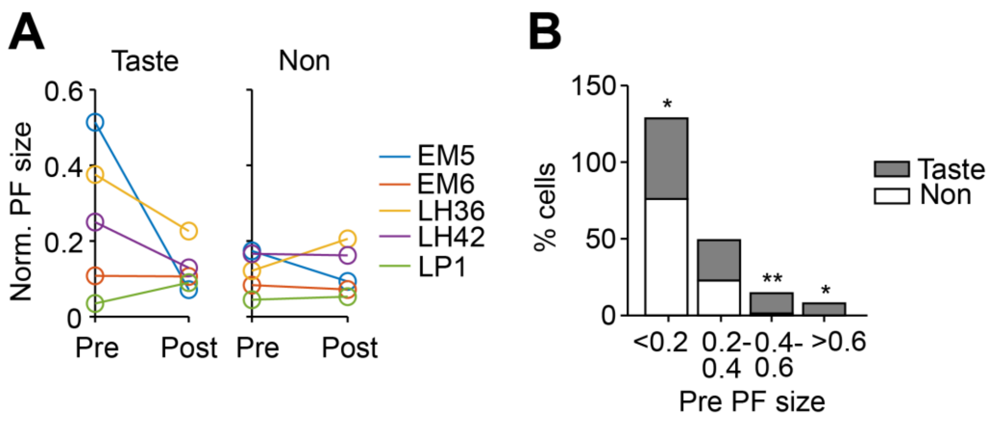


**Figure S2. Taste responsiveness is related to place field size during the Pre session, related to Figure 2.**

***A***, Place field size in **Figure 2B** shown for individual animals. ***B***, Histogram of taste-responsive and non-taste-responsive place field sizes. Stacked bars represent the percentage of cells within each cell type that fell within a given size bin (n*0.2 as a fraction of the track). Cells with smaller place fields were more likely to be non-taste-responsive (PF size < 0.2, $\chi^2$ test, $\chi$ = 6.5, *$p$ = 0.011), while cells with larger place fields were more likely to be taste-responsive (PF size = 0.4-0.6, $\chi$ = 7.5, **$p$ = 0.0063; PF size > 0.6, $\chi$ = 6.4, *$p$ = 0.011).

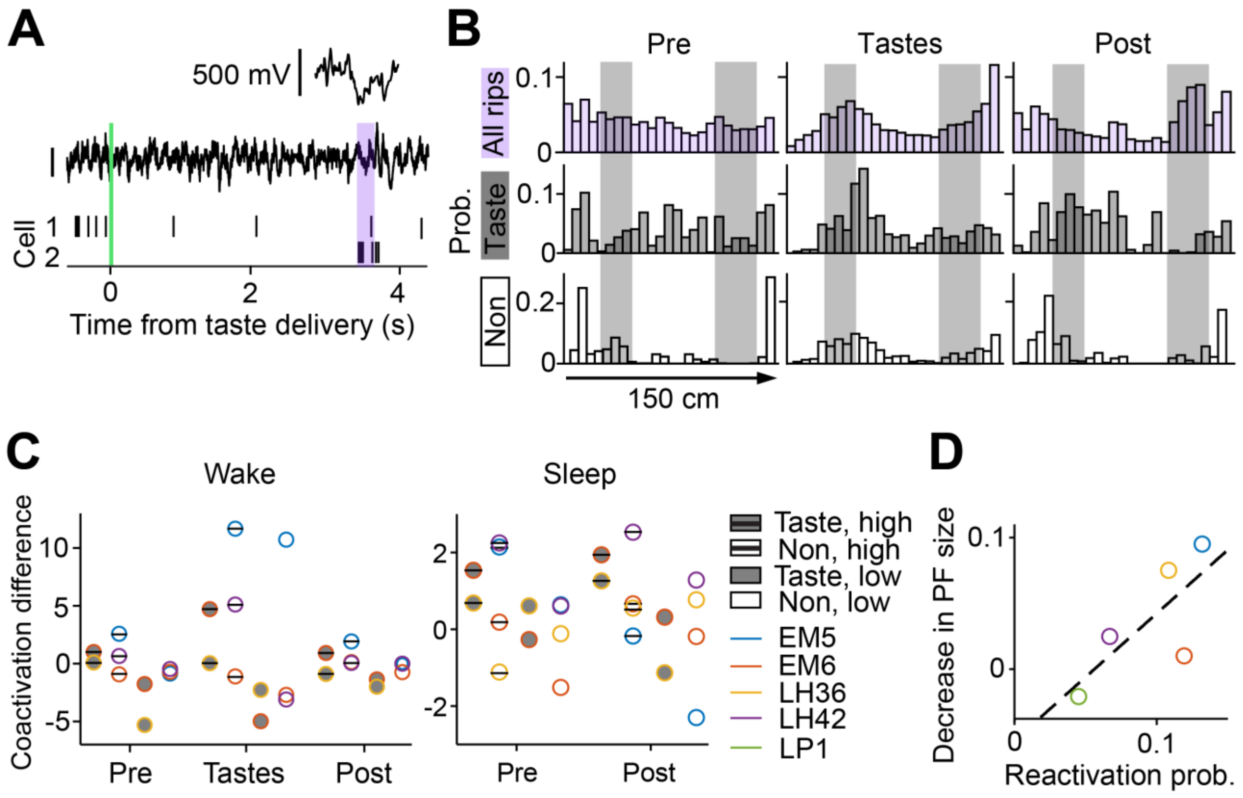


**Figure S3. SWR co-activation and distribution of significant events along the track, related to Figure 4.**

***A***, Example of one co-activation event following a taste trial, with saccharin delivery denoted by the vertical green line. Spikes from two taste-responsive cells are shown in black, with spikes that occurred during the same SWR inside the purple bar. SWRs were detected using the simultaneously recorded EEG filtered at 150-250 Hz (black traces above raster plot). The top trace shows the SWR in higher magnification. ***B***, Probability of SWRs and significant co-activation events for taste- and non-taste-responsive cell pairs at each location on the track during awake running sessions. ***C***, Mean co-activation differences from **Figure 4B** shown for individual animals (each column corresponds to a bar on the main plot). For each epoch, only animals with sufficient numbers of cell pairs are shown. ***D***, Mean reactivation probability vs. decrease in place field (PF) size from **Figure 4C** shown for individual animals.

| Animal | CA1 cells | | | | | |
|---|---|---|---|---|---|---|
| | All | Pyr | Int | Taste-responsive | Pyr | Int |
| EM5 | 35 | 26 | 9 | 7 | 1 | 6 |
| LH36 | 31 | 28 | 3 | 23 | 19 | 4 |
| EM6 | 35 | 29 | 6 | 17 | 11 | 6 |
| LP1 | 11 | 10 | 1 | 4 | 3 | 1 |
| LH42 | 31 | 25 | 6 | 8 | 4 | 4 |
| Total | 143 | 118 | 25 | 59 | 38 | 21 |

**Table S1. Cell distribution across animals, related to Figure 1.**

Summary of the number of taste-responsive and total CA1 cells recorded from each animal. Only cells meeting the inclusion criteria (see STAR Methods) are reported. Putative pyramidal cells (Pyr) and interneurons (Int) were identified on the basis of firing rate and spike width parameters. Neurons were classified as “taste-responsive” if they exhibited responses to taste presence or identity.
